# Photoprotective mechanisms in *Elysia* species hosting *Acetabularia* chloroplasts shed light on host-donor compatibility in photosynthetic sea slugs

**DOI:** 10.1101/2024.02.02.578635

**Authors:** Luca Morelli, Vesa Havurinne, Diana Madeira, Patrícia Martins, Paulo Cartaxana, Sónia Cruz

## Abstract

Sacoglossa sea slugs have garnered attention due to their ability to retain intracellular functional chloroplasts from algae, while degrading other algal cell components. While protective mechanisms that limit oxidative damage under excessive light are well documented in plants and algae, the photoprotective strategies employed by these photosynthetic sea slugs remain unresolved. Species within the genus *Elysia* are known to retain chloroplasts from various algal sources, but the extent to which the metabolic processes from the donor algae can be sustained by the sea slugs is unclear. By comparing their responses to high light conditions through kinetic analyses, molecular techniques, and biochemical assays, this study highlights significant differences between two photosynthetic *Elysia* species with chloroplasts derived from the green alga *Acetabularia acetabulum*. Notably, *Elysia timida* displayed remarkable tolerance to high light stress and sophisticated photoprotective mechanisms such as an active xanthophyll cycle, efficient D1 protein recycling, accumulation of heat-shock proteins and α-tocopherol. In contrast, *Elysia crispata* exhibited absence or limitations in these photoprotective strategies. Our findings emphasize the intricate relationship between the host animal and the stolen chloroplasts, highlighting different capacities to protect the photosynthetic organelle from oxidative damage.

## 1 Introduction

Photosynthesis, a vital process converting light energy to chemical energy, is generally associated to plants and algae. However, sacoglossan sea slugs are intriguing examples of “solar-powered” animals, retaining functional chloroplasts through a process named kleptoplasty (Kawaguti & Yamasu, 1965; Greene, 1970; Greene & Muscatine, 1972; Cruz & Cartaxana, 2022). The supply of photosynthesis-derived metabolites from the stolen chloroplasts (named kleptoplasts once inside the animal tissues) can sustain sea slug weight during periods of food shortage or support reproductive output (e.g., Casalduero & Muniain, 2008; Cartaxana et al., 2021).

However, the relationship between phototrophic organisms and light is intricate. Excessive light generates reactive oxygen species (ROS) in the chloroplasts, causing damage to DNA, lipids, proteins, and other molecules of the cell. Various stressors, including salinity, drought, temperature extremes, and heavy metal exposure exacerbate the production of ROS in photosynthetic organisms, increasing oxidative damage, particularly under high-light conditions (Müller & Munné-Bosch, 2021). Light induced damage to photosystem II (PSII), termed photoinhibition, is largely caused by ROS, singlet oxygen in particular, reacting with proteins of the thylakoid membranes (Vass, 2011; Tyystjärvi, 2013; Nishiyama and Murata, 2014). PSII in photosynthetic organisms consists of various subunits, with some encoded in the chloroplast and others in the nucleus. The reaction centre complex, critical for charge separation, includes plastid-encoded proteins like D1, D2, PsbI, and the α and β subunits of Cyt b559 (PsbE and PsbF, respectively). On the lumenal side of the thylakoid membrane, the oxygen evolving complex comprises three nuclear-encoded proteins (PsbO, PsbP, and PsbQ) (Suorsa et al., 2014; Järvi et al., 2015). Photoinhibition of PSII is inevitable even under low-light, but instead of a complete resynthesis of all the subunits to counteract the damage, only the D1 protein of a damaged PSII is *de novo* synthesised, while most of the other subunits are recycled. This PSII repair cycle is a complex process mediated by multiple auxiliary proteins like kinases, proteases, chaperones, and translocases. In plants, most of these proteins are nucleus encoded (Tyystjärvi & Aro, 1996; Järvi et al. 2015; Theis and Schroda, 2016).

In response to high-light exposure, plants and algae employ xanthophyll cycles (XCs), which involve the reversible conversion of xanthophyll pigments from an epoxidized to a de-epoxidized state. This transformation induces a conformational change in light-harvesting complexes, enhancing the dissipation of excess excitation energy as heat and protecting the photosynthetic apparatus (Latowski et al., 2011; Goss & Jakob, 2010). Additionally, under these conditions, the yield of chlorophyll fluorescence emission is linked to the amount of de-epoxidized pigment in the xanthophyll pool (Frank et al., 2000). The most prevalent type of XC in higher plants and in multiple algae species involves the conversion of violaxanthin (Vx) to zeaxanthin (Zx) via the intermediate antheraxanthin (Ax) under high light conditions, catalysed by the enzyme violaxanthin de-epoxidase (VDE). The process is then reversed by the enzyme zeaxanthin epoxidase (ZEP) under lower irradiances (Latowski et al., 2011).

The XCs contribute to non-photochemical chlorophyll fluorescence quenching (NPQ), a collective term for multiple photoprotective processes occurring in photosynthetic membranes (Demmig-Adams & Adams, 1996). The primary NPQ mechanism affected by the XCs is known as qE, a rapid and reversible process dependent on the irradiance-induced proton gradient across the thylakoid membrane, resulting in a reduction in luminal pH (ΔpH). In land plants, PsbS protein serves as the sensor detecting these changes and initiating qE quenching within PSII antenna complexes (Dall’Osto et al., 2005). Based on data mainly from the unicellular green alga *Chlamydomonas reinhardtii,* the lumenal pH changes are predictably sensed by the protonation of specific stress related light harvesting complex proteins (LHCSRs), mainly LHCSR3, in green algae (Peers et al., 2009). Protonation activates the qE quenching activity of LHCSR3, possibly by allowing it to act as the quenching site by itself, or by causing LHCSR3 to quench excess excitation energy of other light harvesting complexes, like LHCII, via protein-protein interactions (Bonente et al., 2011; Ballottari et al., 2016; Girolomoni et al., 2019; Cazzaniga et al., 2020).

Different mechanisms contributing to NPQ are traditionally identified based on kinetic analyses of their induction and relaxation during and after high light exposure, respectively. qE type quenching is remarkably swift and typically relaxes within 5 min after light removal, following the dissipation of ΔpH (the trigger for qE) that takes only 10 to 20 s in the dark (Zaks et al., 2012). An NPQ component with a relaxation time of approximately 5 to 20 min is associated with state transitions and termed qT. This process involves the reversible redistribution of excitation energy between PSI and PSII mainly in low to moderate light conditions, although it might also be involved with protection against high-light in green algae (Allorent et al., 2013; Derks et al., 2015). The third component in the classical description of NPQ, qI, is associated with photoinhibition of PSII, with a relaxation time of tens of min to h. The qI quenching site was recently shown to likely reside in the D1 protein of a damaged PSII, and the relaxation of qI is linked to D1 degradation by FtsH protease in *C. reinhardtii* (Nawrocki et al., 2021). An additional component, named qZ, was identified based on the correlation between the slow (tens of min) formation and reconversion of Zx and the chlorophyll fluorescence relaxation. qZ is a PsbS- and ΔpH-independent quenching process unrelated to qI and qT. It develops over 10 to 20 min, likely representing a quenching process in the PSII antenna at xanthophyll binding sites that convert to Zx more slowly than the dynamic Vx to Zx conversion (Nilkens et al., 2010). These mechanisms are well-documented in higher plants. However, in some algae species, acidification of the thylakoid lumen activates the XC, but qE appears to be absent (Garcia-Mendoza et al., 2011).

Conversely, in certain unicellular green algae, XC does not contribute to qE, despite qE being a significant part of their NPQ (Quaas et al., 2015). On another hand, in some Bryopsidales algae, XC is entirely absent, and NPQ slowly increases under illumination, without relaxing during the recovery phase (Christa et al., 2017; Giovagnetti et al., 2018).

The diversity of photoprotective mechanisms becomes crucial when considering sea slugs that integrate foreign chloroplasts, but do not preserve algal nuclear genes needed for the encoding of most proteins with a central role in chloroplast maintenance. For example, kleptoplasts from *Elysia timida* and *Elysia chlorotica* exhibit rapid and reversible NPQ, along with a functional XC, mirroring their presence in the respective algal prey (*Acetabularia acetabulum* and *Vaucheria litorea*, respectively) (Cruz et al., 2015; Cartaxana et al., 2019). However, when XC-competent chloroplasts from *Chaetomorpha* sp. were integrated by *Elysia viridis*, a sea slug known for its prolonged kleptoplast retention when fed with appropriate algae, this mechanism was not retained (Morelli et al., 2023; Rauch et al., 2018). The full spectrum of photoprotective mechanisms in photosynthetic sea slugs remains to be fully explored, especially as different combinations of animal hosts and algal chloroplast donors can result in a variety of maintenance processes.

*Elysia crispata* is a polyphagous sea slug that feeds on a multitude of different algae in the wild, and it was recently shown to be able to incorporate long-term functional chloroplasts also from the XC-competent alga *A. acetabulum* (Cartaxana et al., 2023). We hypothesize that chloroplasts originating from a shared algal donor may trigger distinct metabolic adaptations, contingent upon the specific animal species hosting them as kleptoplasts. Thus, this study compares the photoprotective mechanisms employed by *E. crispata* fed with *A. acetabulum* to those of the monophagous *E. timida* that only feeds on and incorporates chloroplasts from *A. acetabulum*. Our research sheds light on the importance of host-donor compatibility for long-term kleptoplasty in photosynthetic sea slugs.

## 2 Materials and Methods

### 2.1 Animals and algae collection and maintenance

*E. crispata* sea slugs originally collected in Florida (USA) and purchased from TMC Iberia (Lisbon, Portugal) were reared in 150 L recirculated life support systems (LSS) operated with artificial seawater (ASW) at 25 °C and a salinity of 35 ppt. The population of *E. timida* used in this study originates from the Mediterranean (Elba, Italy, 42.7782° N, 10.1927° E) and has been routinely cultured in the laboratory for years, as described by Havurinne and Tyystjärvi (2020). Macroalgae were cultivated separately to feed sea slugs. The green alga *Bryopsis plumosa*, acquired from Kobe University Macroalgal Culture Collection (KU-0990, KUMACC, Japan), was grown in 2 L flasks with f/2 medium lacking Na_2_SiO_3_ and constant aeration at 20 °C and an irradiance of 60-80 μmol photons m^−2^ s^−1^ provided by LED lamps (Valoya 35 W, spectrum NS12). The green alga *A. acetabulum* (strain DI1 originally isolated by Diedrik Menzel) was cultivated essentially as described by Havurinne and Tyystjärvi (2020) in 3-20 L transparent plastic boxes filled with f/2 medium lacking Na_2_SiO_3_, without aeration under the same lamps as *B. plumosa*. The photon scalar irradiance was set to 40 μmol photons m^−2^ s^−1^ for *A. acetabulum*. For both algae, the photoperiod was the same as the one used for the sea slugs. For this work, *E. crispata* specimens raised and fed with *B. plumosa* were continuously fed with *A. acetabulum* for 10 days to induce kleptoplast change as observed and described by Cartaxana et al. (2023).

### 2.2 Experimental protocols

#### 2.2.1 Light-stress recovery

To assess the photoprotective capabilities of both algae and animals, samples underwent a light stress and recovery (LSR) protocol involving sequential phases: 15 min of darkness (D), followed by 20 min of high-light (HL, 1200 μmol photons m^−2^ s^−1^), and finishing with 40 min of recovery under low-light (LL_Rec_, 40 μmol photons m^−2^ s^−1^). The blue light LED of an Imaging-PAM fluorometer (Mini version, Heinz Walz GmbH) was used as the light source for the LSR protocol. The number or samples used in every single experiment is indicated in the figure legends.

#### 2.2.2 Chemical pre-treatments

For the lincomycin treatment, sea slugs and algae were submerged in a 10 mM solution of lincomycin hydrochloride (Lin) in ASW and kept in darkness for 12 h before experiments. As an alternative to the direct treatment, *E. timida* individuals were also fed lincomycin-treated algae for one week to ensure a consistent supply of affected chloroplasts. This was done after a 10-day starvation period, followed by daily feeding with 100 mg of *A. acetabulum* filaments pre-treated with lincomycin, discarding the previous day’s filaments. Post-treatment, the animals were utilized in experiments. For 1,4-dithiothreitol (DTT) treatments, animals and algae were immersed in a 10 mM DTT solution in darkness for 2 h before starting the experiments. This DTT solution was also used to prepare the agar embedding solution for the sea slugs, as described in Section 2.3.

### 2.3 Photosynthetic measurements

Variable fluorescence measurements were carried out at room temperature by using an Imaging-PAM fluorometer (Mini version, Heinz Walz GmbH). To avoid animal movements, the animals were embedded in a 0.2% agar solution in ASW before starting the measurements. The agar solution was cooled to roughly 37 °C on ice prior to pouring it on top of the sea slugs to minimize the heat stress of the animals and to ensure fast fixation. Effective quantum yield of photosystem II (ΦPSII) was measured as ΔF/Fm’, where ΔF corresponds to Fm’-F (the maximum minus the minimum fluorescence of light-exposed organisms). Maximum quantum yield of PSII (Fv/Fm), was calculated as (Fm-Fo)/Fm, where Fm and Fo are, respectively, the maximum and the minimum fluorescence of dark-adapted samples. For simplicity, only the term ΦPSII is used in the text and figures, and Fv/Fm is referred to as ΦPSII after dark acclimation. The NPQ kinetics were obtained by recording chlorophyll *a* variable fluorescence values in samples exposed to the LSR protocol (see section 2.2.1). NPQ was calculated as (Fm-Fm’)/Fm’.

### 2.4 Pigments and α-tocopherol analysis

Pigment analysis was performed as described by Cruz et al. (2014). Briefly, sea slugs and algae were sampled from the respective experimental protocol and immediately frozen in liquid nitrogen. Samples were freeze-dried, powdered by using a fine metal rod and pigments extracted in 95% cold buffered methanol (2% ammonium acetate) by adding the buffer, sonicating the samples for 2 min, and then incubating them at −20°C for a period of approximately 20 min. The extracts were subsequently cleared of debris by filtration through 0.2 μm Fisherbrand™ PTFE membrane filters and then injected into an HPLC system (Shimadzu, Kyoto, Japan) equipped with photodiode array (SPD-M20A) and fluorescence (RF-20A) detectors. Pigments were identified from absorbance spectra and retention times and concentrations were calculated using the peak areas in the photodiode array detector, in comparison with pure crystalline standards (DHI, Hørsolm, Denmark). The activity of the XC and the sequential de-epoxidation of the pigments Vx to Ax and Zx, was estimated by calculating the de-epoxidation state (DES) as: DES=([Zx]+0.5×[Ax])/([Zx]+[Ax]+[Vx]). To analyse α-tocopherol, identification and concentration calculations were conducted by observing the retention time and peak areas in the fluorescence detector (excitation: 295 nm; emission 330 nm) in comparison to a pure α-tocopherol standard (Sigma-Aldrich, St. Louis, MO, USA).

### 2.5 Chloroplast molecular identification

Total genomic DNA was extracted from sea slugs using the DNeasy Blood and Tissue DNA extraction kit (Qiagen, Hilden, Germany), following the manufacturer’s protocol. Partial regions of the plastid *rbcL* (∼600 bp) gene was amplified by using a nested PCR approach. Amplification of the *rbcL* gene fragments was performed in a Veriti 96-well Thermal Cycler (Applied Biosystems) with the primer pair rbcLF (5′-AAAGCNGGKGTWAAAGAYTA −3′) and rbcLR (5′-CCAWCGCATARAWGGTTGHGA −3′) (Pierce et al., 2006). Reaction mixtures (25 µL) contained 12.5 μL DreamTaq™ PCR Master Mix (Fisher Scientifc), 0.1 µM of each primer and 1 µL of template DNA. PCR for amplification of *rbcL* gene was performed employing the following thermal cycling conditions: an initial denaturation at 95 °C for 1 min, followed by 35 cycles of denaturation at 94 °C for 45 s, annealing at 47 °C for 45 s. An extension at 72 °C for 90 s was also performed, followed by a final extension at 72 °C for 10 min. Amplicons of the first PCR were used as template for a second amplification with identical cycling parameters but 25 denaturation cycles using again the primers rbcLF and rbcLR. PCR-amplified fragments were sent to a certified laboratory (STAB VIDA, http://www.stabvida.com/) for purification and Sanger sequencing. rbcL nucleotide sequences were sequenced in a single direction. To confirm species identification, a BLAST search (Basic Local Alignment Search Tool, BLAST, National Center for Biotechnology Information, NCBI) was finally performed. The sequences generated in this study were deposited in GenBank® under accession numbers PP114081-PP114090.

### 2.6 Protein analysis

Total protein extraction was carried out following a modified version of the protocol described by Shanmugabalaji et al. (2013). Approximately 5 mg of freeze-dried material was ground with 300 µL of lysis buffer (100 mM Tris-HCl pH 7.7, 2% SDS, 0.1% protease inhibitor (Sigma), 100 mM N-acetyl-L-cystein) vigorously shaken for 1 min, and then incubated at 37 °C for 30 min with constant shaking. The supernatant was collected after centrifuging the samples for 15 min at 10,000 *g* and the protein amount was calculated by mixing the samples with the Bradford reagent and measuring the optical absorbance at 595 nm. Proteins were then precipitated by adding 4 volumes of acetone and by incubating the samples at –20 °C for 30 min. After centrifuging the samples for 15 min at 10,000 *g* at 4 °C the upper phase was removed and the pellet washed with 500 µL of 80% acetone, dried and resuspended in Laemmli Sample Buffer (Bio-Rad), prepared according to the manufacturer instructions to obtain a final protein concentration of 2 µg µL^-1^. SDS-PAGE separation was performed by loading 20 µg of protein samples previously boiled at 95 °C for 10 min on a 12-4% (separating– stacking) polyacrylamide gel. Western blot was performed by using a Trans-Blot Turbo transfer system and following the manufacturer instructions (Standard SD program: 30 min, 25 V constant). Nitrocellulose membranes were stained with Ponceau solution before antibody incubation to check the success of the transfer process. To probe the blot, a primary antibody recognising PsbA (D1) protein was used (Agrisera, Sweden). The secondary antibody was anti-rabbit IgG conjugated with horseradish peroxidase (Agrisera). Chemiluminescence was detected using SuperSignal ECL substrate (Thermo Fischer Scientific) and developed using a Chemidoc XRS imaging system (Biorad).

The chaperone protein content (HSP70/HSC70) was assessed to examine the protein quality control mechanisms in response to oxidative stress. Initially, 10 µL of the homogenate supernatant was diluted in 250 µL of PBS. Following this, 50 µL of the diluted sample was dispensed into 96-well microplates (Nunc-Roskilde, Denmark) and allowed to incubate overnight at 4 °C. After 24 h, the microplates underwent washing with PBS containing 0.05% Tween-20 (40%, Sigma-Aldrich). Subsequently, 200 µL of a blocking solution (1% bovine serum albumin, Sigma-Aldrich) was added to each well and left to incubate at room temperature for 2 h. Following another round of microplate washing, 50 µL of a 5 mg mL^-1^ solution of primary antibody (anti-HSP70/HSC70, Acris, Rockville, MD, US) was introduced to each well and incubated at 37 °C for 90 min. After removing non-linked antibodies via further washing, alkaline phosphatase-conjugated anti-mouse IgG (Fab specific, Sigma-Aldrich) was utilized as a secondary antibody. This involved adding 50 µL of a 1 mg mL^-1^ solution to each well and incubating the microplates for an additional 90 min at 37 °C. Following three additional washing steps, 100 µL of substrate (SIGMAFASTTM p-nitrophenyl phosphate tablets, Sigma-Aldrich) was dispensed into each well and allowed to incubate for 10–30 min at room temperature. Subsequently, 50 µL of stop solution (3 M NaOH, ≥98%, Sigma-Aldrich) was introduced to each well, and the absorbance was read at 405 nm using a 96-well microplate reader (Asys UVM 340, Biochrom, US). The concentration of HSP70/HSC70 in the samples was calculated based on an absorbance curve derived from serial dilutions (ranging between 0 and 2 mg mL^-1^) of purified HSP70 active protein (Acris). Results were expressed relative to the protein content of the samples, determined by Bradford assay (Bradford, 1976).

## 3 Results

### 3.1 Photoprotection in *E. timida* and *E. crispata* fed with *A. acetabulum*

A molecular verification based on DNA barcoding was employed to confirm the origin of the kleptoplasts within *E. crispata* specimens that had been raised in the laboratory on *B. plumosa* (Ec-Bp) and then nourished with *A. acetabulum* continuously for ten consecutive days (Ec-Aa). The *rbcL* gene sequences from the sampled sea slugs were compared with sequences available in GenBank using the BLAST-n search tool to identify matching sequences. The sequences from Ec-Bp shared an average of 99.90% of similarity with *B. plumosa* and *Bryopsis hypnoides* (100% query with 100% similarity to one *rbcL* sequence and 100% query with 99.59% similarity to one *rbcL* sequence available on GenBank®, respectively). In contrast, the sequences from Ec-Aa and *E. timida* displayed 100% similarity with *A. acetabulum* (100% query with 100% similarity to one *rbcL* sequence available on GenBank®) (Supplementary Figure S1).

Following the consumption of *A. acetabulum*, the coloration of *E. crispata* specimens became akin to that of *E. timida*, a result of both sharing the same chloroplast donor (Figure 1A). While ΦPSII after dark acclimation did not exhibit differences between *E. crispata* and *A. acetabulum*, it was significantly higher in *E. timida* (0.649±0.010, 0.652±0.023, and 0.682±0.021, respectively). Under HL, the ΦPSII approached near-zero values in both sea slugs and alga. After a 40-min low-light recovery (LL_Rec_) phase, both *E. timida* and *A. acetabulum* regained a significant portion of their ΦPSII, with values of 0.442±0.009 and 0.406±0.043, respectively, showing no statistically significant differences. However, *E. timida* displayed a notably faster increase in the recovery of photosynthetic efficiency. In fact, 5 min after transition to low-light, *E. timida* showed a ΦPSII value of 0.321±0.021, while *A. acetabulum* exhibited a value of 0.151±0.036. In contrast to especially *E. timida*, *E. crispata* only reached ΦPSII values of 0.237±0.022 at the end of LL_Rec_, showing a slower recovery compared to the other specimens (Figure 1B). Regarding NPQ, *E. crispata* achieved a NPQ value of 3.00±0.17 during HL but reduced it only to 2.43±0.23 during LL_Rec_. In *E. timida* and *A. acetabulum*, on the other hand, NPQ reached 2.50±0.41 and 2.37±0.14 during HL and subsequently decreased to 0.90±0.09 and 0.67±0.18, respectively, during LL_Rec_ (Figure 1C).

**Figure 1.**
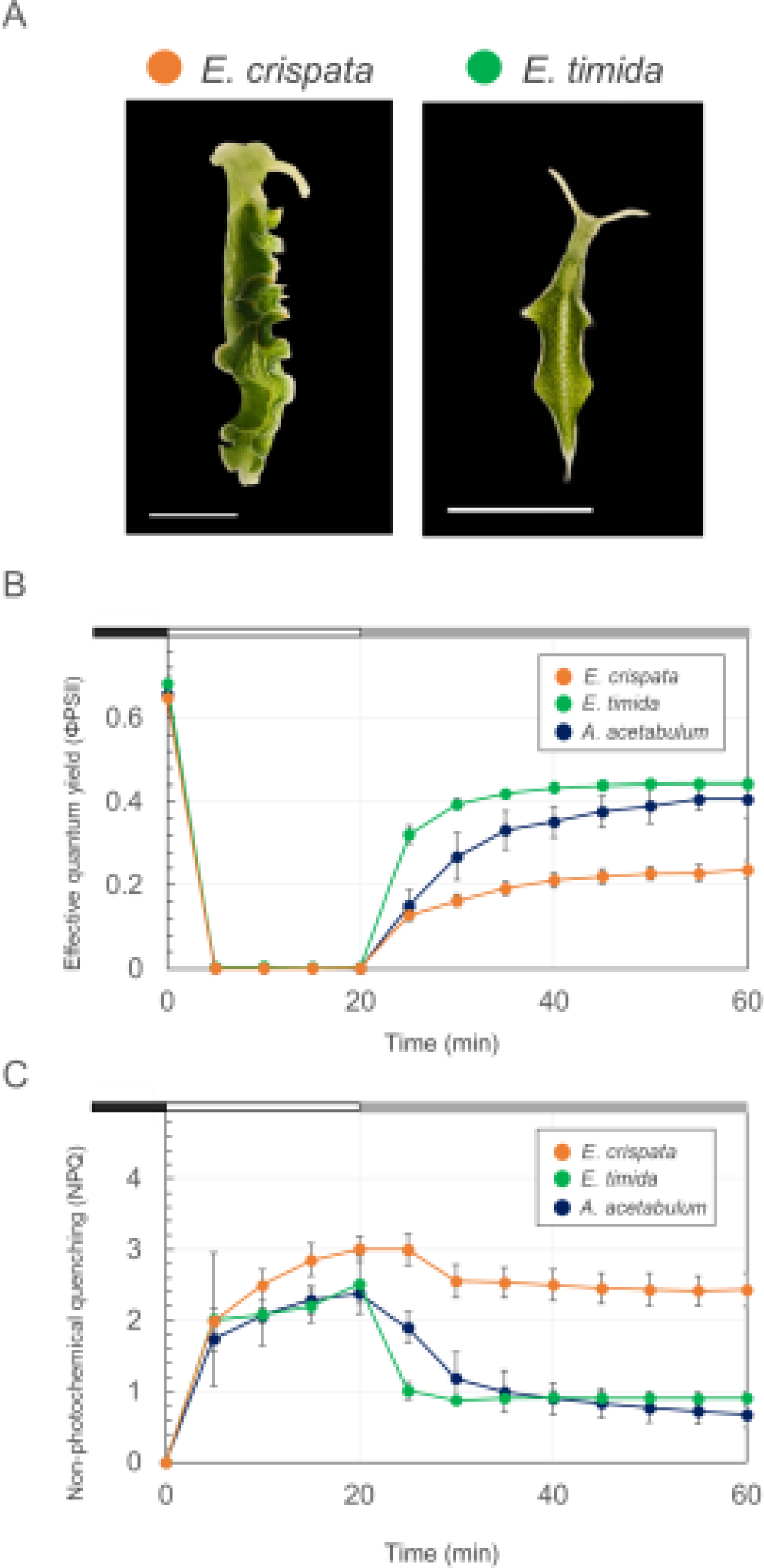
Light stress and recovery in *Elysia crispata*, *Elysia timida*, and *Acetabularia acetabulum*. **(A)** Representative images of the two sea slug species fed with the same chloroplast donor (*A. acetabulum*), with a scale bar representing 5 mm. **(B)** Effective quantum yield (ΦPSII) measured in *E. crispata*, *E. timida*, and *A. acetabulum* during a light stress and recovery experiment. The chart highlights different protocol phases: black bar for dark acclimation (15 min), white bar for high-light stress (1200 μmol photons m^-2^ s^-1^ for 20 min), and grey bar for low-light recovery (40 μmol photons m^-2^ s^-1^ for 40 min), displaying mean and standard deviation (n=5). **(C)** Non-photochemical quenching (NPQ) kinetics under the same conditions as described for (B).

Subsequently, the XC operation was examined in *E. crispata*, *E. timida* and *A. acetabulum*, in the three phases of the LSR protocol. Neither Ax nor Zx were detected in any of the samples before HL exposure, while dark-adapted. *E. timida* and *A. acetabulum* showed an active XC, with Vx converting to Zx under HL and back to Vx during LL_Rec_ (Figure 2A). This led to significantly higher de-epoxidation state (DES) values under HL compared to the ones found in LL_Rec_, in these samples (Figure 2B). In contrast, *E. crispata*, when exposed to HL, accumulated Zx which did not revert to Vx during the recovery phase. This resulted in a sustained DES value without significant change between HL and LL_Rec_ (Figures 2A, 2B).

**Figure 2.**
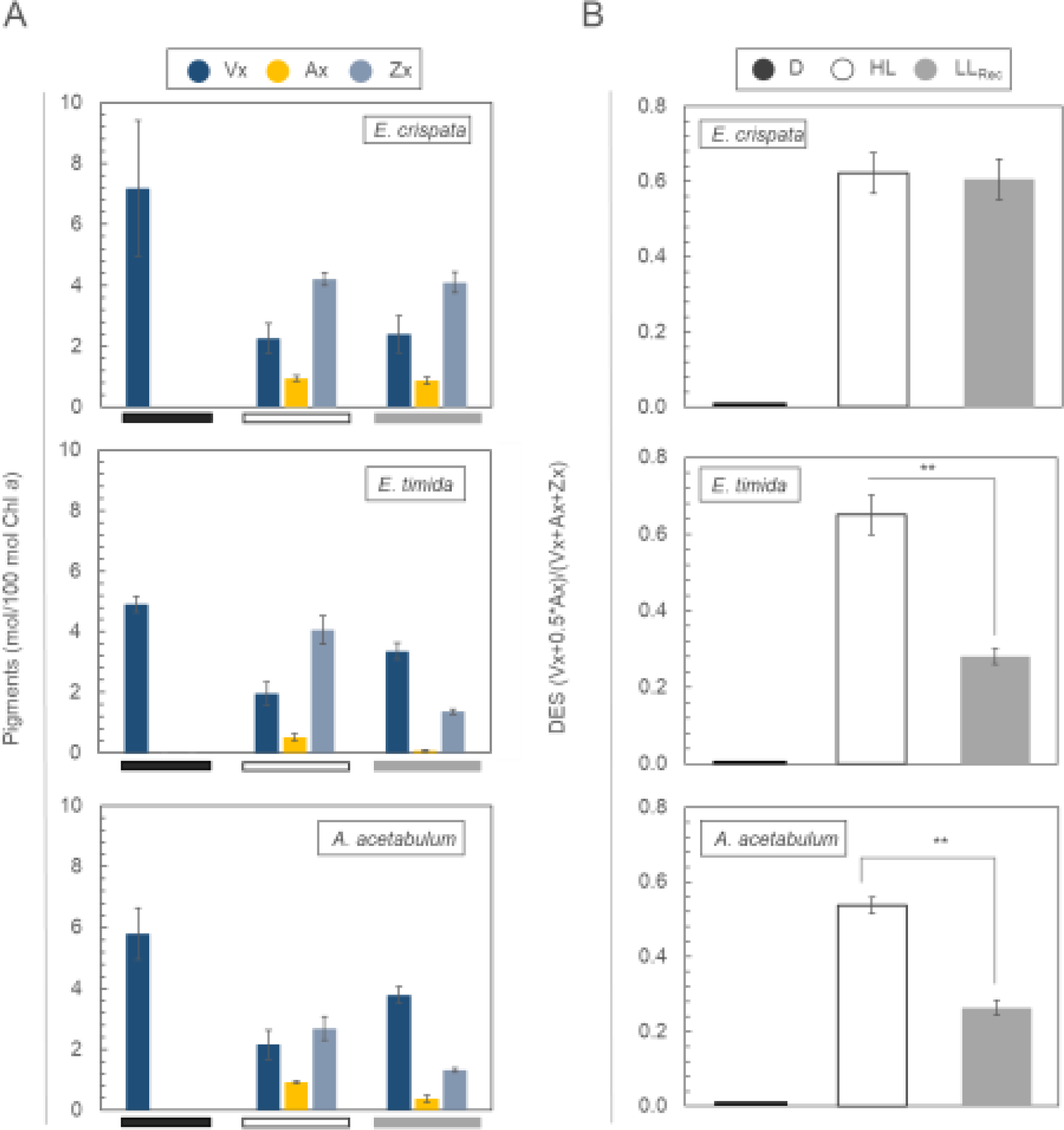
Operation of the xanthophyll cycle in *Elysia crispata, Elysia timida,* and *Acetabularia acetabulum.* **(A)** Levels of the single xanthophylls expressed as mol of pigment relative to 100 mol of chlorophyll (Chl) *a* during a light stress and recovery experiment; black bar: dark-acclimated for 15 min; white bar: high-light stress, 1200 μmol photons m^-2^ s^-1^ for 20 min; and grey bar: low-light recovery, 40 μmol photons m^-2^ s^-1^ for 40 min. **(B)** de-epoxidation state (DES) expressed as (Zx+0.5*Ax)/(Vx+Ax+Zx) in samples subjected to a light stress and recovery protocol as described for (A). D, dark; HL, high-light; LL_Rec_, low-light recovery; Vx, violaxanthin; Ax, antheraxanthin; Zx, zeaxanthin. Data corresponds to mean and standard deviation (n=4). Asterisks mark statistically significant differences between HL and LL_Rec_ phases (t-test, ** p < 0.01).

The D1 protein, a constituent of the core of PSII, was detected by Western blot analysis in *E. crispata*, *E. timida*, and *A. acetabulum*. This protein was observed to undergo dynamic degradation under HL and regeneration in LL_Rec_ as a photoprotective response, in both sea slugs species and *A. acetabulum* (Figure 3). Although resynthesis of damage D1 was observed in all three samples under LL_Rec_, *E. timida* seemed more efficient in recovering the original D1 pool following the HL stress.

**Figure 3.**
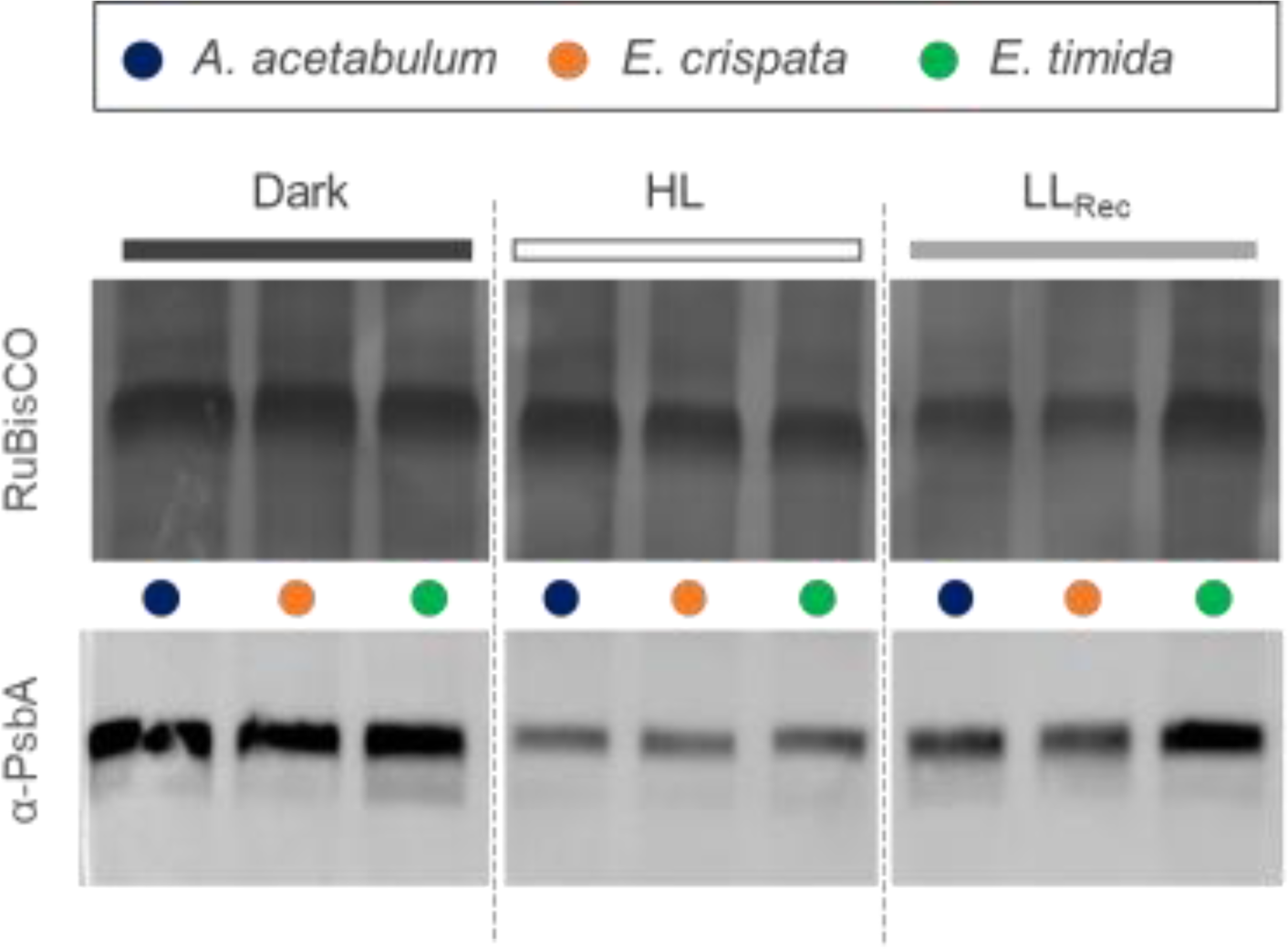
Western blot showing the levels of D1 protein in *Acetabularia acetabulum*, *Elysia crispata*, and *Elysia timida* during a light stress and recovery experiment; Dark: dark-acclimated for 15 min; HL: high-light stress, 1200 μmol photons m^-2^ s^-1^ for 20 min; and LL_Rec_: low-light recovery, 40 μmol photons m^-2^ s^-1^ for 40 min. The loading control is the RuBisCO band found in the same samples and stained by using Ponceau solution. The arrangement of the samples, coming from the same membrane, has been adjusted for consistency with the figures in the main text.

### 3.2 Effect of chloroplast impairment on photoprotective mechanisms

Lincomycin (Lin) treatment, lasting 12 h, was used to inhibit chloroplast protein synthesis in the sea slugs. Lin-treated samples displayed a significantly lower ΦPSII after dark acclimation (Fv/Fm) compared to untreated samples (Figure 4A). Lin-treated *E. crispata* and *A. acetabulum* samples showed reduced photosynthetic yield recovery compared to their untreated counterparts (Figures 4B, 1B). Specifically, ΦPSII values of the Lin-treated samples after LL_Rec_ were 0.107±0.017 for *E. crispata* and 0.136±0.033 for *A. acetabulum* (Figure 4B), as opposed to the higher values (0.237±0.022 and 0.406±0.043) observed in the untreated samples (Figure 1B). On the other hand, Lin-treated *E. timida* recovered most of its photosynthetic activity, reaching 0.366±0.022 after recovery, near the value showed by untreated specimens (0.442±0.009) (Figures 4B, 1B). NPQ kinetics mirrored this recovery. Lin-treated *E. crispata* and *A. acetabulum* reached NPQ values of 1.97±0.206 and 2.27±0.207 after HL but dissipated this slowly, resulting in NPQ values of 1.37±0.165 and 1.69±0.041, respectively, after LL_Rec_ (Figure 4B). Notably, Lin-treated *A. acetabulum* exhibited NPQ kinetics like untreated *E. crispata* (Figure 1C). Conversely, *E. timida* effectively dissipated NPQ from 1.73±0.220 in HL to 0.596±0.169 after LL_Rec_ in the presence of lincomycin. XC operation was influenced by chloroplast protein synthesis inhibition as *E. crispata* and *A. acetabulum* increased Zx under HL but failed to revert it to Vx during LL_Rec_, showing unvarying DES values (Figure 4C). On the contrary, *E. timida* exhibited a functional XC in the presence of lincomycin, with DES decreasing from 0.456±0.017 (HL) to 0.321±0.009 after low-light recovery. These results were confirmed in *E. timida* specimens fed Lin-treated algae (Supplementary Figure S2). Despite this different administration method, *E. timida* exhibited an unchanged ability to maintain an active XC and to recover photosynthetic activity after high-light stress.

**Figure 4.**
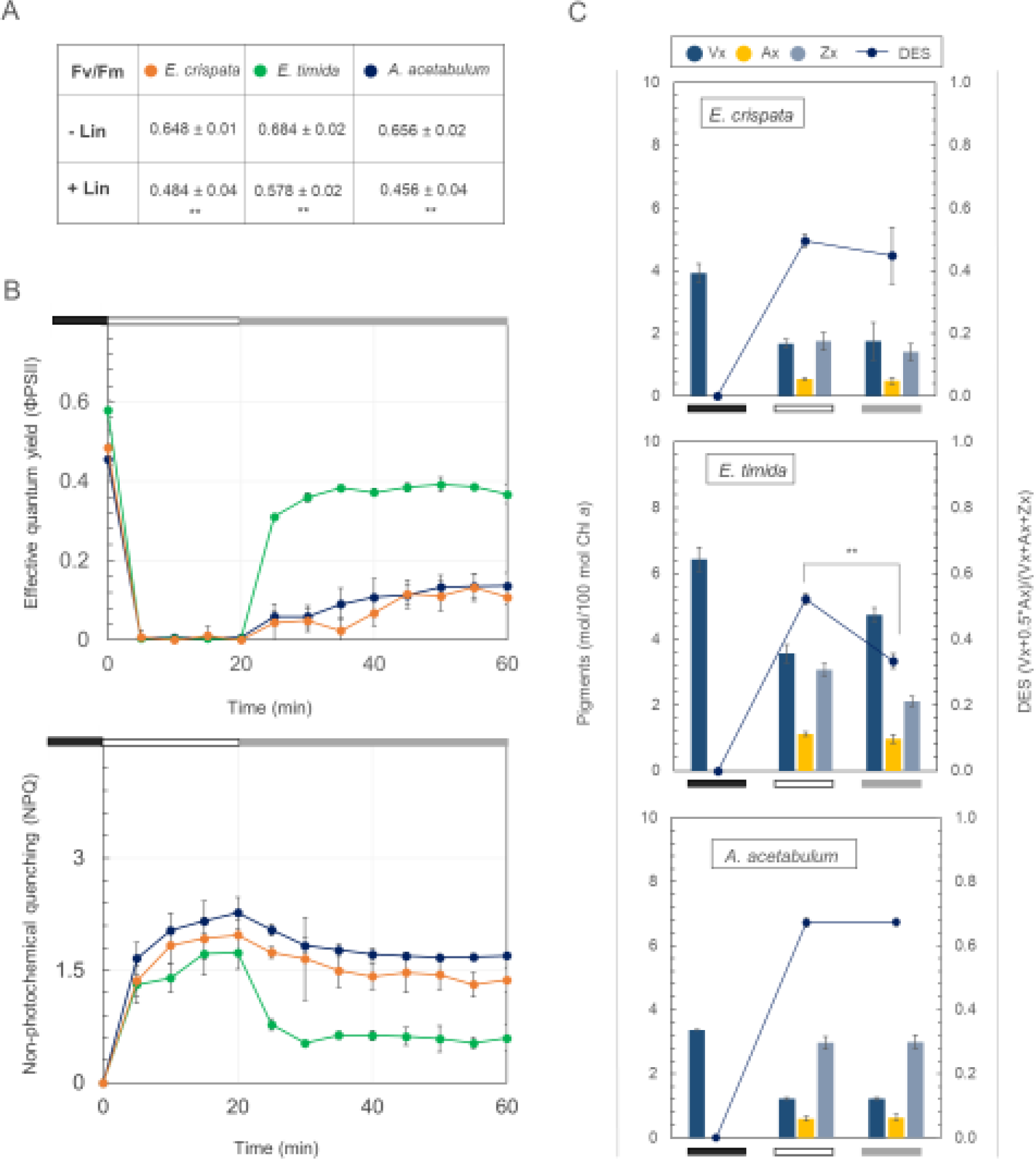
Effects of chloroplast protein synthesis inhibition on a light stress and recovery experiment and respective xanthophyll cycle operation in *Elysia crispata*, *Elysia timida,* and *Acetabularia acetabulum*. **(A)** Variation of maximum quantum yield (Fv/Fm) in *E. crispata*, *E. timida*, and *A. acetabulum* in the absence (-Lin) and presence (+Lin) of the inhibitor lincomycin hydrochloride. Asterisks mark statistically significant differences between –Lin and +Lin (t-test, ** p < 0.01). **(B)** Effective quantum yield (ΦPSII) and non-photochemical quenching (NPQ) measured in lincomycin treated samples during a light stress and recovery experiment. The chart highlights different protocol phases: black bar for dark acclimation (12 h), white bar for high-light stress (1200 μmol photons m^-2^ s^-1^ for 20 min), and grey bar for low-light recovery (40 μmol photons m^-2^ s^-1^ for 40 min), displaying mean and standard deviation (n=5). **(C)** Operation of the xanthophyll cycle measured at the end of each protocol phase described in (B). The bar plots show the levels of the single xanthophylls expressed as mol of pigment relative to 100 mol of chlorophyll (Chl) *a*. The lines show the de-epoxidation state (DES) expressed as (Zx+0.5*Ax)/(Vx+Ax+Zx) in samples subjected to the light stress and recovery protocol; Vx, violaxanthin; Ax, antheraxanthin; Zx, zeaxanthin. Data corresponds to mean and standard deviation (n=4). Asterisks mark statistically significant differences between the high-light and the recovery in low-light (t-test, ** p < 0.01).

DTT application effectively inhibited the XC in all samples (*E. crispata*, *E. timida*, and *A. acetabulum*), as evidenced by the absence of Ax and Zx under HL, indicating inhibition of the enzyme violaxanthin de-epoxidase (Figure 5A). With a non-functional XC, all samples exhibited near null photosynthetic activity (0.0225±0.0175, 0.0267±0.0128, and 0.0212±0.0166 for *E. crispata*, *E. timida*, and *A. acetabulum,* respectively) and built up limited NPQ levels (barely reaching the value of 1) during HL. During recovery under low-light, *E. crispata* and *A. acetabulum* showed no significant change in ΦPSII and NPQ values compared to high-light. Contrastingly, *E. timida* showed a significant increase in ΦPSII (from 0.056±0.013 to 0.210±0.019) and a decrease in NPQ (from 0.89±0.074 to 0.70±0.009) (Figure 5B).

**Figure 5.**
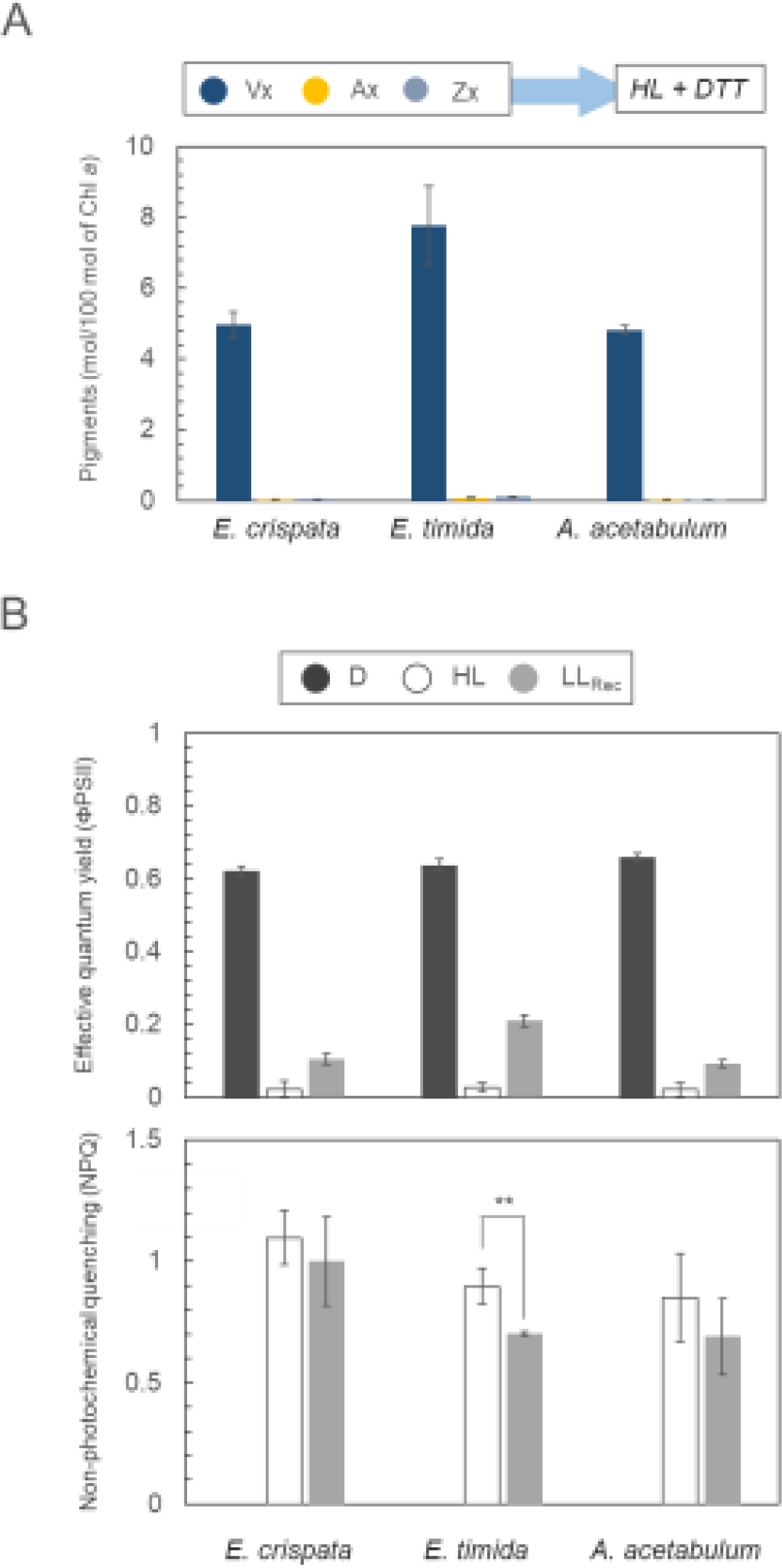
Effects of the inhibition of the xanthophyll cycle by 1,4-dithiothreitol (DTT) in *Elysia crispata*, *Elyisa timida,* and *Acetabularia acetabulum*. **(A)** Levels of the single xanthophylls expressed as mol of pigment relative to 100 mol of chlorophyll (Chl) *a* observed in the DTT-treated samples during high-light exposure. **(B)** Effective quantum yield (ΦPSII) and non-photochemical quenching (NPQ) in samples treated with DTT and exposed to the light stress and recovery experiment. D, dark for 2 h; HL, 1200 μmol photons m^-2^ s^-1^ for 20 min; LL_Rec_, 40 μmol photons m^-2^ s^-1^ for 40 min. Data corresponds to mean and standard deviation (n=4). Asterisks mark statistically significant differences between the high-light and the recovery in low-light (t-test, ** p < 0.01).

### 3.3 Accumulation of heat-shock proteins and antioxidants

The protein HSP70, involved in stabilizing other proteins in response to environmental stress, was found to accumulate at significantly higher levels in *E. timida* compared to *E. crispata* during the LSR protocol (Figure 6A). Remarkably, HSP70 levels continued to rise in *E. timida* from 22.38±1.27 µg mg^-1^ of total proteins under HL to 45.27±8.27 µg mg^-1^ at the end of the LL_Rec_ phase. A similar, but more dynamic pattern emerged in the accumulation of α-tocopherol, a metabolite with potent antioxidant properties. While the levels of the compound in *E. crispata* and *A. acetabulum* remained constant during HL and after transferring the samples to recovery conditions, α-tocopherol significantly increased in *E. timida* upon HL exposure (i.e., going from 24.26±2.04 pg mg^-1^ DW in D to 48.69±6.59 pg mg^-1^ DW in HL) subsequently decreasing (31.51±5.12 pg mg^-1^ DW) during LL_Rec_ (Figure 6B). Interestingly, among the three analysed samples, *A. acetabulum* was the one showing lowest α-tocopherol levels.

**Figure 6.**
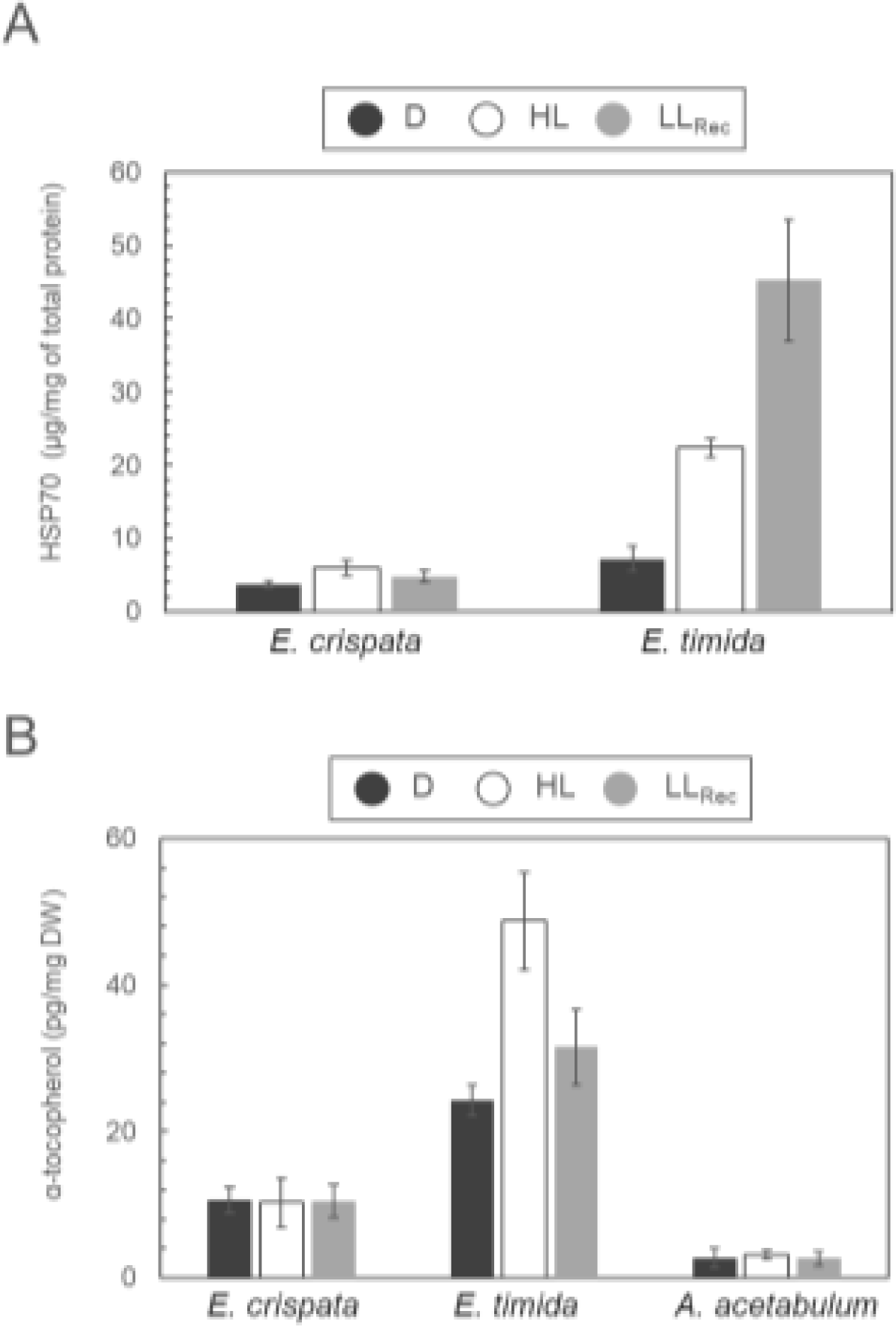
Accumulation of heat-shock proteins and antioxidants. **(A)** Levels of HSP70 protein (μg/mg of total protein) in *E. crispata* and *E. timida* during a light stress and recovery experiment. **(B)** α-tocopherol levels in *E. crispata*, *E. timida*, and *A. acetabulum* (pg/mg DW) in the same conditions. D, dark for 15 min; HL, 1200 μmol photons m^-2^ s^-1^ for 20 min; LL_Rec_, 40 μmol photons m^-2^ s^-1^ for 40 min. Data corresponds to mean and standard deviation (n=4).

## 4 Discussion

In plants and algae, many photoprotection mechanisms are regulated through gene expression, with light intensity and spectrum playing essential roles in controlling chloroplast function, acting through nuclear-encoded receptors and signalling pathways (Duan et al., 2020; Morelli et al., 2021, Pinnola & Bassi, 2018). However, in photosynthetic sea slugs this interplay remains largely mysterious because chloroplasts are the sole algal components integrated into the animal cells, with the nucleus and other cellular components being degraded (Kawaguti & Yamasu, 1965; Greene, 1970; Greene & Muscatine, 1972). To begin unravelling this puzzle, we initiated an analysis involving two distinct sea slug species sharing the same algal donor: *E. crispata* and *E. timida* feeding on the green alga *A. acetabulum*. Previous studies have highlighted the broad dietary preferences of adult *E. crispata*, feeding on over 30 ulvophycean algal species from the Bryopsidales and Dasycladales orders (Middlebrooks et al., 2014, 2019; Vital et al., 2021, 2023). In laboratory conditions, *E. crispata* that normally feed on *B. plumosa* successfully integrated kleptoplasts from *A. acetabulum* in just 10 days (Cartaxana et al., 2023) as confirmed visually (Figure 1A) and now by PCR-based DNA barcoding using the chloroplast-encoded *rbcL* sequence (Supplementary Figure S1).

*A. acetabulum* chloroplasts are known to have a fully functional XC that contributes to the qE component of NPQ in a typical manner, leading to NPQ kinetics where the amplitude of its build-up and relaxation correlates with the rapid changes in ΔpH when exposed to high-light and subsequently put to recover in low-light or darkness (Christa et al., 2017; Zaks et al., 2012; Johnson et al., 2008). Although there were slight differences in especially the NPQ relaxation kinetics, we confirmed that *E. timida* and *A. acetabulum* samples exhibit similar NPQ kinetics (Figure 1), and akin to those found in other algae and sea slugs known for their reversible NPQ and XC, such as *V. litorea* and *E. chlorotica* (Cruz et al., 2015). In contrast, *E. crispata* displayed a slow and limited NPQ relaxation under low irradiance, seemingly devoid of the fast-relaxing qE component, suggesting that some form of sustained quenching dominates the NPQ process more in *E. crispata* than in *E. timida* or *A. acetabulum.* When we quantified the operation of the XC, we observed that *E. crispata* exhibited an increase in Zx content during high-light exposure, which did not decrease during the subsequent low-light recovery period. A similar scenario was reported also in the polyphagous sea slug *E. viridis* that shifted its diet specifically to the XC-competent alga *Chaetomorpha* (Morelli et al., 2023). The continued presence of Zx in *E. crispata* during low-light recovery may suggest a reliance on some form of sustained quenching, with the most likely candidate being qZ, as qZ requires Zx for operation and functions independently of ΔpH changes (Nilkens et al., 2010). *E. crispata*’s inability to revert Zx to Vx could lock the quenching into this slower mechanism, perhaps even providing some benefits purely in terms of photoprotection in highly dynamic light environments.

Multiple studies have established a clear connection between Zeaxanthin Epoxidase (ZEP) protein stability and the degree of photoinhibition. The abundance of ZEP protein and its conversion of Zx to Ax and subsequently to Vx are closely linked to recovery from photoinhibition (Jahns and Miehe, 1996; Verhoeven et al., 1996). Moreover, ZEP activity is progressively reduced in response to decreasing PSII activity during high-light stress, and hydrogen peroxide produced by PSI in high-light has been suggested to be the main ROS inhibiting ZEP activity in plants (Holzmann et al., 2022; Kress and Jahns, 2017; Reinhold et al., 2008). Recent research has hinted at the susceptibility of the ZEP enzyme to more controlled redox regulation involving the thioredoxin and glutathione networks in *Arabidopsis thaliana* chloroplasts (Naranjo et al., 2016). We postulate that, under normal conditions, *A. acetabulum* can deal with the electron pressure caused by the transition to and during high-light by using carbon fixation, alternative electron acceptors such as flavodiiron proteins downstream of PSI, and the native antioxidant systems of the alga without major ROS bursts. Both *E. timida* and *E. crispata* have been shown to be capable of carbon fixation via their chloroplasts (Trench, 1969; de Vries et al., 2015; Cartaxana et al., 2021), suggesting that carbon fixation also works as an electron sink in both species. *E. timida* has also been shown to retain the flavodiiron activity of *A. acetabulum* chloroplasts, although the activity seems weaker in *E. timida* compared to the alga according to spectroscopic P700 oxidation measurements (Havurinne and Tyystjärvi, 2020). The flavodiiron activity was not measured in *E. crispata*, but if it is compromised as in *E. timida*, the chloroplast flavodiiron proteins in these sea slugs might not effectively prevent hydrogen peroxide accumulation, unlike in *A. acetabulum*. This could lead to detrimental hydrogen peroxide levels in high-light, potentially inhibiting ZEP specifically in *E. crispata*. *E. timida*, with a more dynamic metabolic response to high-light, including α-tocopherol synthesis, should be better equipped to mitigate the inhibition of ZEP by ROS (Figure 6).

When treated with lincomycin, an inhibitor of chloroplast protein synthesis, *A. acetabulum* exhibited an NPQ phenotype akin to that of *E. crispata*. In contrast, *E. timida*, while experiencing a reduction in maximum photosynthetic activity, sustained a functional xanthophyll cycle and NPQ relaxation even under conditions of protein synthesis inhibition (Figure 4 and Supplemental Figure S2). As ZEP is encoded in the nucleus, its synthesis should remain largely unaffected by lincomycin. Therefore, ZEP’s loss of function in the presence of lincomycin likely stems from the inhibition of thylakoidal protein turnover and, consequently, the repair of PSII as observed in other systems like *Arabidopsis* (Bethmann et al., 2019).

The unique ability of *E. timida* to maintain a functional xanthophyll cycle post-lincomycin treatment, and the capacity of this species of progressively recovering photosynthetic activity while unable to perform XC because of DTT inhibition (Figure 5) raises the hypothesis that this species may employ alternative mechanisms to ensure enzyme stability and to tackle potentially dangerous ROS levels. For example, *E. timida* demonstrated a notable increase in the production and responsiveness of HSP70 upon exposure to high-light stress. HSP70 proteins, belonging to the heat shock protein family, target hydrophobic stretches within unfolded or misfolded polypeptides, aiding in proper folding, preventing the formation of inactive or toxic aggregates, and facilitating the degradation of irreversibly damaged proteins (Mayer et al., 2005). The upregulated expression of HSPs stands as a fundamental cellular defence mechanism against protein misfolding and damage provoked by environmental stressors (Tomanek, 2008; Pulido et al., 2017). This responsive cascade is prominently observed across various photosynthetic organisms. Furthermore, it has been postulated that plastidial proteases, notably FtsH and Deg, play an instrumental role in the detoxification of protein aggregates under stress conditions. This includes their involvement in the proteolysis of the D1 protein, especially under high-intensity light stress. In this situation, in *Arabidopsis*, cytosolic HSP70 has been proposed to undergo upregulation and to interact with the FtsH1 subunit, not only to bolster the stability of the FtsH1 subunit but also to ensure its efficient translocation into the chloroplasts to participate in chloroplast functionality support (Shen et al., 2007). Considering the significant increase in HSP70 production during light stress and recovery in *E. timida*, we suggest that in this species an increase of this protein may have a protective role in preserving plastidial components or proteins from degradation, possibly contributing in this way to preserve photoprotective mechanisms for longer. The exceptionally high HSP levels in *E. timida*, when compared to *E. crispata* suggest that *E. timida* is in a class of its own in terms of adaptability. A differential response to environmental challenges in terms of HSP70 production had already been described for the photosynthetic sea slugs *E. viridis* and *E. crispata* (Dionísio et al., 2018), and in the case of the *Aiptasia*-algal symbioses where this class of proteins participates in the maintenance of the photosynthetic activity of the algae symbiont in case of heat stress (Petrou et al., 2021).

The dynamic of α-tocopherol accumulation can also reflect this capacity of *E. timida* to respond to environmental challenges in a fast an efficient way. In plants, α-tocopherol is promptly synthesized in response to high-light to tackle ROS production at the thylakoidal level and reduce photodamage (Munné-Bosh, 2005). The same molecule also acts as a powerful intracellular signalling molecule controlling redox homeostasis (Munné-Bosh, 2007). One cannot entirely dismiss the possibility that, under environmental stress, healthy and non-aging *E. timida* kleptoplasts produce substantial amounts of antioxidants. These molecules could not only mitigate damage by detoxifying ROS but also signal the sea slug about internal threats, triggering chaperone and support peptide production to alleviate stress impacts. Such synergistic responses are likely in the highly specialized *E. timida*, due to the evolutionary compatibility between the animal and the chloroplasts of its sole food source, *A. acetabulum*. However, this intricate interaction is less probable in cases like *E. crispata*, where this specific and unique association is not as well established (Figure 7).

**Figure 7.**
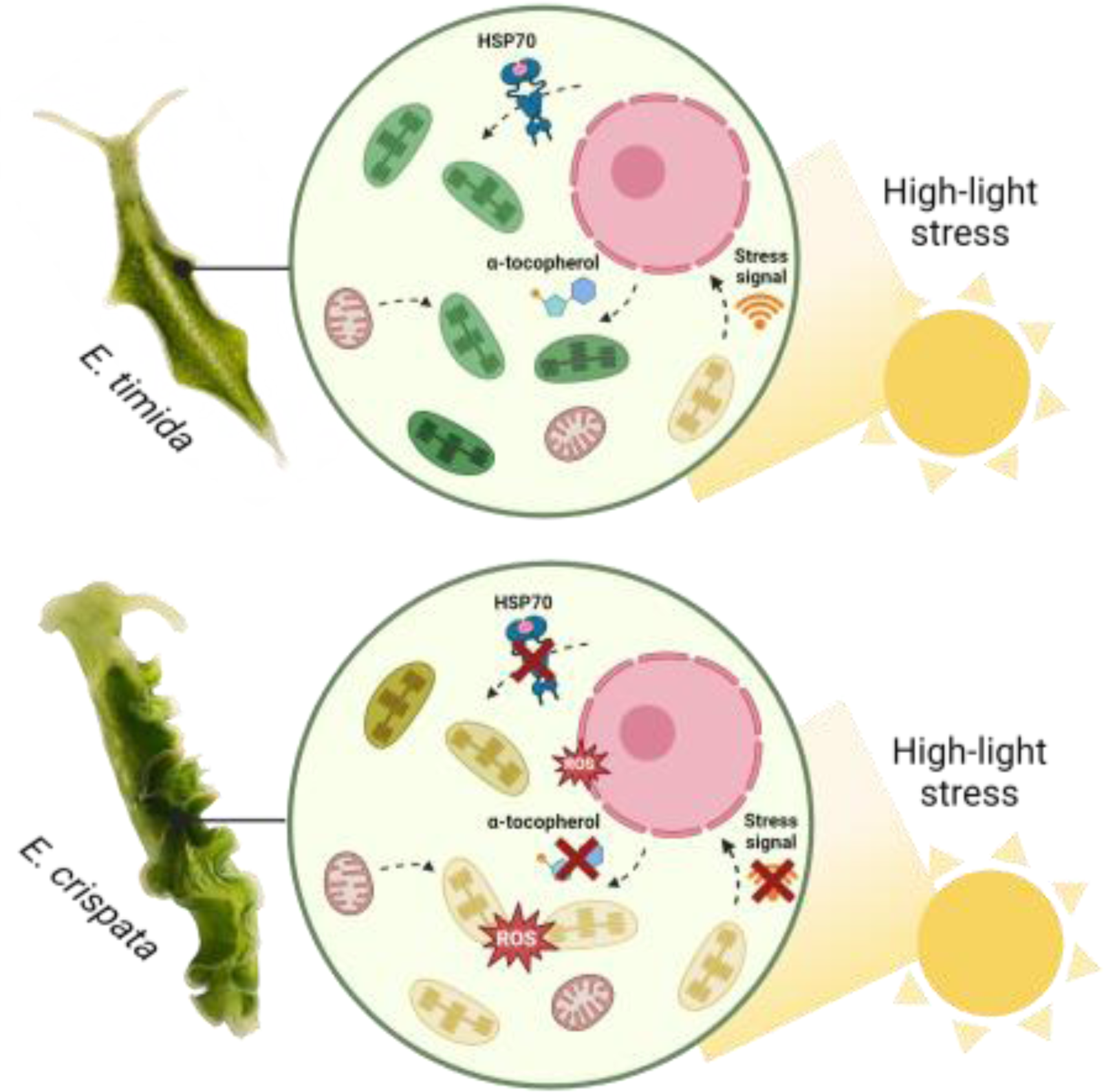
Schematic representation of *Acetabularia acetabulum* chloroplast maintenance in *Elysia* species. In *E. timida*, high-light stress triggers stress signals from the chloroplast which prompt photoprotective mechanisms (xanthophyll cycle and protein D1 degradation and resynthesis) and the synthesis of antioxidant products and chaperones (α-tocopherol and HSP70 protein) that help maintain chloroplast function. In *E. crispata*, these mechanisms are generally absent, putatively because the association with the algal chloroplast has not the same degree of specificity. The graphic was generated using BioRender.com.

## 5 Conclusion

Our study sheds light on the photoprotective strategies of photosynthetic sea slugs harbouring algal chloroplasts. While their specific strategies are yet to be fully categorized, we found remarkable differences in the response of different sea slug species to high-light stress, even when sharing the same chloroplast donor. *E. timida* shows high tolerance to high-light stress and the conservation of advanced photoprotective mechanisms under several conditions possibly related to alternative strategies for enzyme and structural protein stability of the sea slug (e.g., HSP70 and tocopherol production). *E. crispata*, despite hosting kleptoplasts coming from the same alga cannot maintain most photoprotective mechanisms. These findings open the door to further investigations into the unique adaptations of kleptoplastidic sea slugs, highlighting that the compatibility between the animal host and algal donor extends far beyond the ability to ingest and integrate the chloroplasts.

## Author contributions

L.M.: conceptualization, investigation, methodology, visualization, writing— original draft, review and editing; V.H.: resources, methodology, writing—review and editing; D.M.: resources, methodology writing—review and editing; P.M.: methodology, formal analysis, writing—review and editing; P.C.: conceptualization, methodology, visualization, writing—review and editing; S.C.: conceptualization, funding acquisition, methodology, project administration, resources, supervision, writing—review and editing.

## Acknowledgments

We thank Maria Inês Silva for technical support in algae and sea slugs maintenance. We also thank Roc Guma Andreu and Marc Canadell (Fundaciò La Pedrera – International stay Joves I Ciencia, Barcelona, Spain) for helping in α-tocopherol quantification.

## Data availability statement

The data that support the findings of this study are available from the corresponding author upon reasonable request.

## Funding

This work was supported by the European Research Council (ERC) under the European Union’s Horizon 2020 research and innovation programme, grant agreement no. 949880 to S.C. (DOI: 10.3030/949880), and by Fundação para a Ciência e Tecnologia, grants 2020.03278.CEECIND to S.C. (DOI:10.54499/2020.03278.CEECIND/CP1589/CT0012), CEECIND/01434/2018 to P.C. (DOI: 10.54499/CEECIND/01434/2018/CP1559/CT0003), CEECIND/00153/2022 to D.M., and UIDP/50017/2020 + UIDB/50017/2020 + LA/P/0094/2020 to CESAM.

---

**Supplementary Figure S1.**
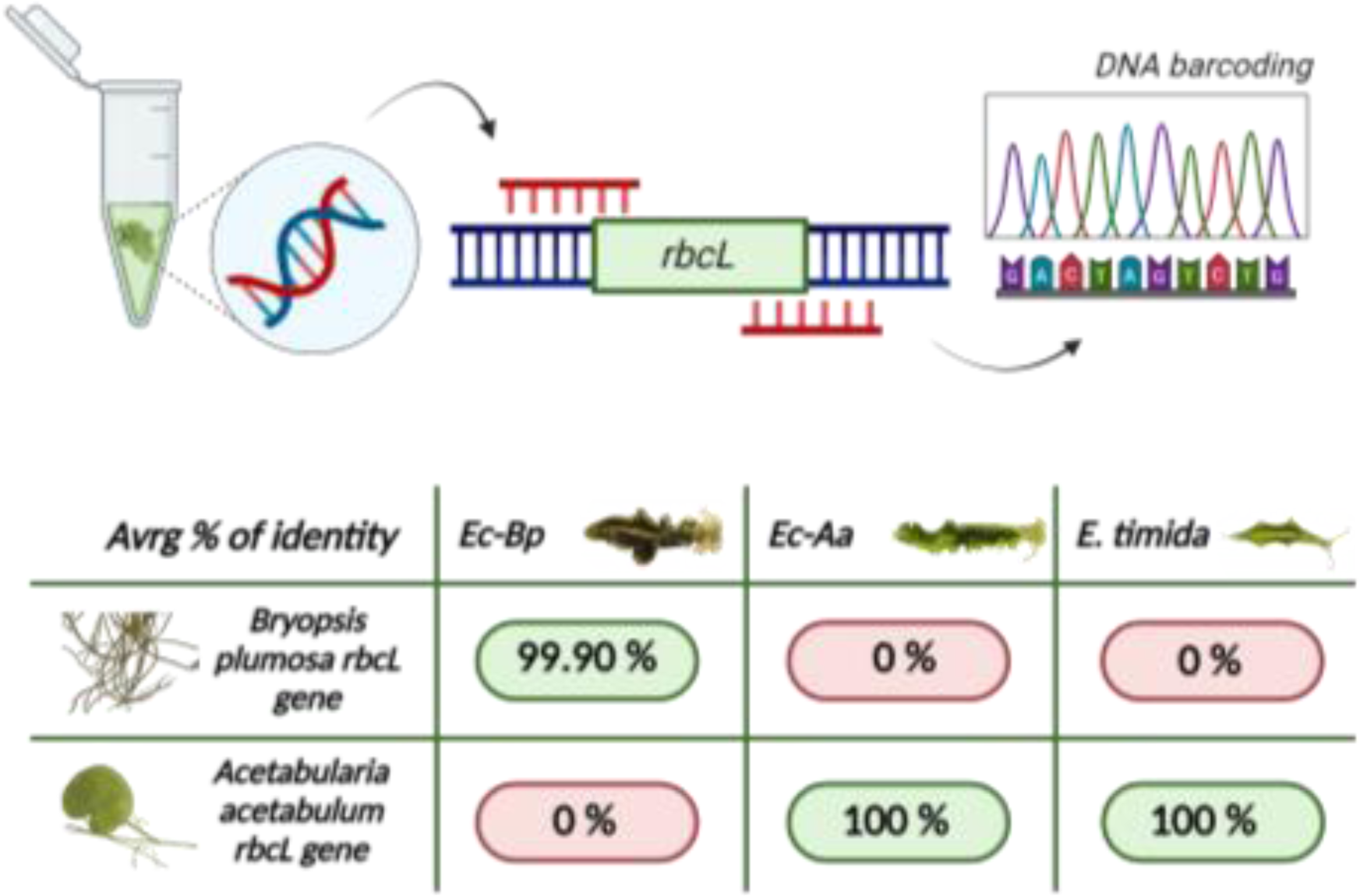
Average percentage of identity between the rbcL gene sequences from sea slugs and those in the database from the corresponding algal food. Ec-Bp: *Elysia crispata* fed *Bryopsis plumosa*; Ec-Aa: *Elysia crispata* fed *Acetabularia acetabulum*; *E. timida*: *Elysia timida*. The graphic was generated using BioRender.com.

**Supplementary Figure S2.**
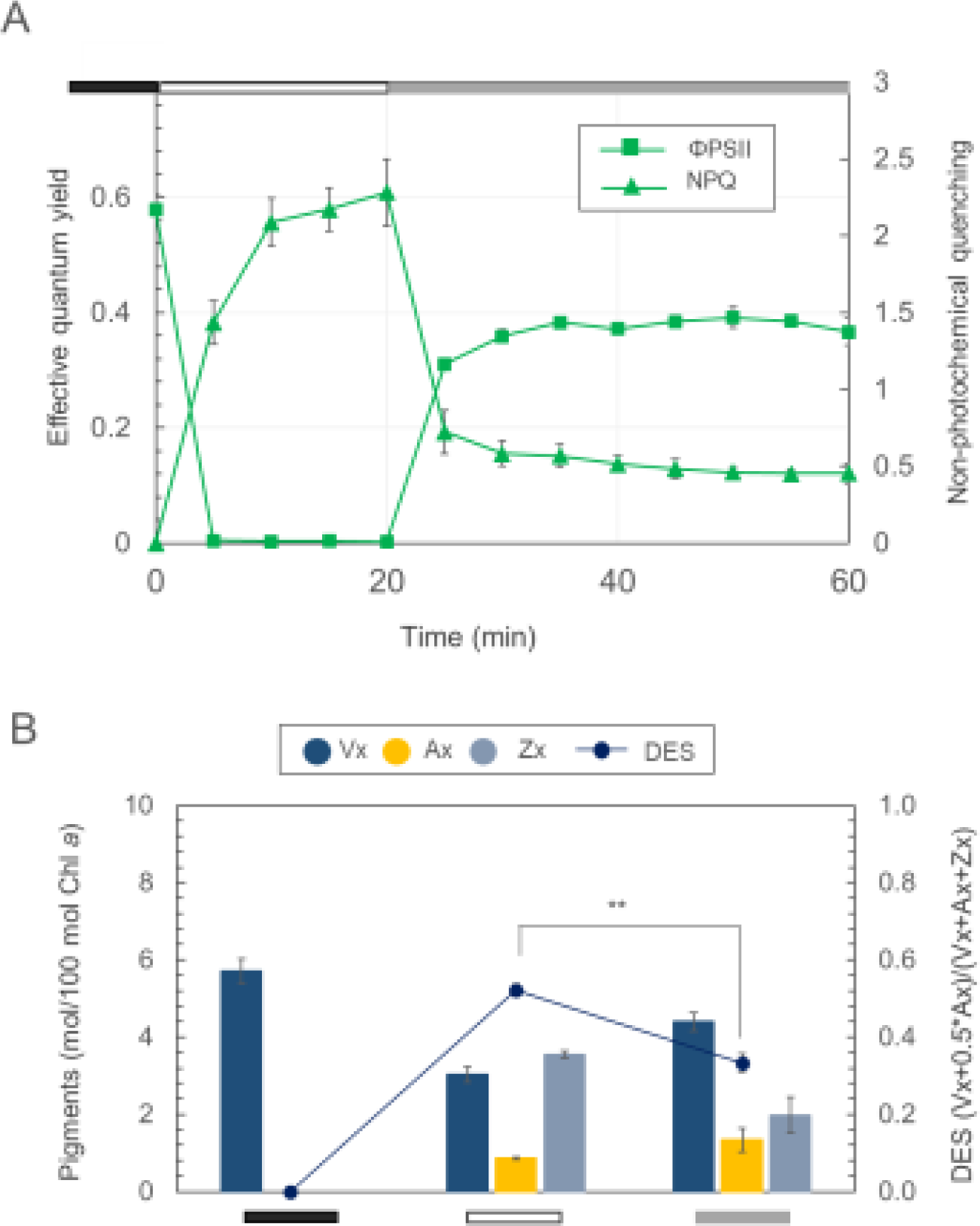
Light stress-recovery and operation of the xanthophyll cycle in *Elysia timida* fed for one week with lincomycin treated *Acetabularia acetabulum* as an alternative to direct chemical treatment of the animals **(A)** Variation of effective quantum yield (ΦPSII) and non-photochemical quenching (NPQ) during a light stress and recovery experiment. The chart highlights different protocol phases: black bar for dark acclimation (15 min), white bar for light stress (1200 μmol photons m^-2^ s^-1^ for 20 min), and grey bar for low light recovery (40 μmol photons m^-2^ s^-1^ for 40 min), displaying mean and standard deviation (n=5). **(B)** Operation of the xanthophyll cycle by showing the levels of the single xanthophylls expressed as mol of pigment relative to 100 mol of chlorophyll (Chl) *a*. The line shows the de-epoxidation state (DES) expressed as (Zx+0.5*Ax)/(Vx+Ax+Zx); Vx, violaxanthin; Ax, antheraxanthin; Zx, zeaxanthin. Data corresponds to mean and standard deviation (n=4). Asterisks mark statistically significant differences between the high light stress and the recovery in low light (t-test, ** p < 0.01).

## Notes

### Competing Interest Statement

The authors have declared no competing interest.

### Summary of Updates

Figures size has been uploaded to solve the problem of the bioRxiv stamper appearing in the middle of each figure.

